# Structural basis for RNA slicing by a plant Argonaute

**DOI:** 10.1101/2022.07.15.500266

**Authors:** Yao Xiao, Shintaro Maeda, Takanori Otomo, Ian J. MacRae

## Abstract

Argonaute (AGO) proteins use small RNAs to recognize transcripts targeted for silencing in plants and animals. Many AGOs possess an endoribonuclease activity termed RNA slicing, which catalyzes rapid turnover of target RNAs. The nuclease activity of *Thermus thermophilus* AGO (TtAgo)^1,2^ is often used as a model for RNA slicing, but how well DNA-guided slicing by this bacterial thermophile resembles RNA-guided slicing by eukaryotic AGOs is not known. We present cryo-EM structures of the *Arabidopsis thaliana* Argonaute10 (AtAgo10)-guide RNA complex with and without a target RNA representing a slicing substrate. Like TtAgo, AtAgo10 conformation expands in response to target binding. However, the AtAgo10-guide-target complex adopts slicing-competent and -incompetent conformations that are distinct from structures of TtAgo. AtAgo10 slicing activity is licensed by docking target (t) nucleotides t9-t13 into a surface channel containing the AGO endoribonuclease active site. A ß-hairpin, conserved in the L1 domain of eukaryotic AGOs, secures the t9-t13 segment and coordinates t9-t13 docking with extended guide-target pairing, preventing the complex from becoming trapped in slicing-incompetent conformations. Results show the mechanism for achieving RNA slicing in eukaryotes is distinct from that of bacteria and provide insights for controlling small interfering RNA (siRNA) potency.

## Main

Small RNAs, including microRNAs (miRNAs) and small interfering RNAs (siRNAs), are essential regulators of gene expression in plants and animals^3,4^. Small RNAs function as guides for Argonaute (AGO) proteins, directing them to complementary sites in RNAs targeted for silencing. The most broadly conserved silencing mechanism used by AGO proteins is endoribonucleolytic target cleavage, or “RNA slicing”^5^. RNA slicing drives RNA interference (RNAi) in animals^6^ and is central to therapeutic siRNA activity in humans^7^. RNA slicing is also essential in plants, where it is the predominant mechanism used by miRNAs^8^ and is used to trigger the biogenesis of phased siRNAs from target transcripts^9-11^.

The slicing mechanism has been characterized in *Thermus thermophilus* AGO (TtAgo), a DNA-guided bacterial AGO that can cleave both DNA and RNA targets^1,2,12^. TtAgo nucleates guide-target base-pairing using the guide (g) proximal seed region (nucleotides g2–g4, counting from the guide 5’ end) and propagates base-pairing to the guide 3’ end^1,13,14^. Target slicing is licensed by extended guide-target pairing, which induces large structural rearrangements in the TtAgo PIWI domain, forming the endonuclease active site^1,2^.

TtAgo is widely referenced as a model for RNA slicing in eukaryotes^4,5,15-18^. However, it is not known if the RNA guides of eukaryotic AGOs behave like the DNA guides of TtAgo. Indeed, structural and biochemical studies suggest mammalian Ago2 propagates guide-target pairing in a manner distinct from the linear 5’-3’ trajectory proposed for TtAgo^15,19,20^. Moreover, the PIWI domains in established eukaryotic AGO structures contain a preformed endoribonuclease active site^21^, indicating that mechanisms distinct from the structural rearrangements observed in TtAgo are used to license target slicing. Previous attempts to crystallize eukaryotic AGOs in an RNA-slicing conformation have instead captured alternative, non-catalytic conformations associated with miRNA-mediated repression and miRNA-decay^19,20^. Thus, despite its fundamental role in small RNA biology and significance to the development of siRNA therapeutics, steps leading to RNA-guided slicing, including how AGOs achieve a slicing-competent conformation and the mechanisms licensing target cleavage, remain unknown.

### Plant and animal AGOs share a common core structure

To visualize the RNA slicing mechanism without trapping the AGO complex in a particular conformation, we employed cryo-EM and single-particle analysis. We found *Arabidopsis thaliana* Ago10 (AtAgo10) behaves particularly well on cryo-EM grids, allowing us to determine its structure despite its small size (110 kDa). As no structure of a plant AGO-RNA complex has yet been reported, we first determined the structure of the AtAgo10-guide RNA complex to 3.3 Å overall resolution (Fig. 1a, S1, and Table S1). As in human AGO structures, AtAgo10 contains N-terminal (N), Linker-1 (L1), PAZ, Linker-2 (L2), MID, and PIWI domains, which form a bilobed structure with a central cleft that contains the guide RNA. AtAgo10 has an additional β-finger structure, not present in human AGOs, that extends 20 Å from the body of the PIWI domain (Fig. 1b, c). The ß-finger is present in clade I AGOs in *Arabidopsis* and found in representatives from all four plant phyla (Fig. 1d, S2a), indicative of an ancient and conserved function in plants. Apart from the β-finger, the AtAgo10 C*α* backbone aligns to human AGOs with an average RMSD of 1.22 Å and is nearly indistinguishable from human AGO structures in an all-against-all correspondence analysis (Fig. 1e, S2b-c). AtAgo10 also displays the guide RNA seed region (g2–g7) in a pre-organized conformation identical to seed display in human AGOs (Fig. 1f)^22-26^. Thus, AGO-guide RNA complexes from Plantae and Animalia kingdoms share a common and deeply conserved core structure.

**Fig. 1.**
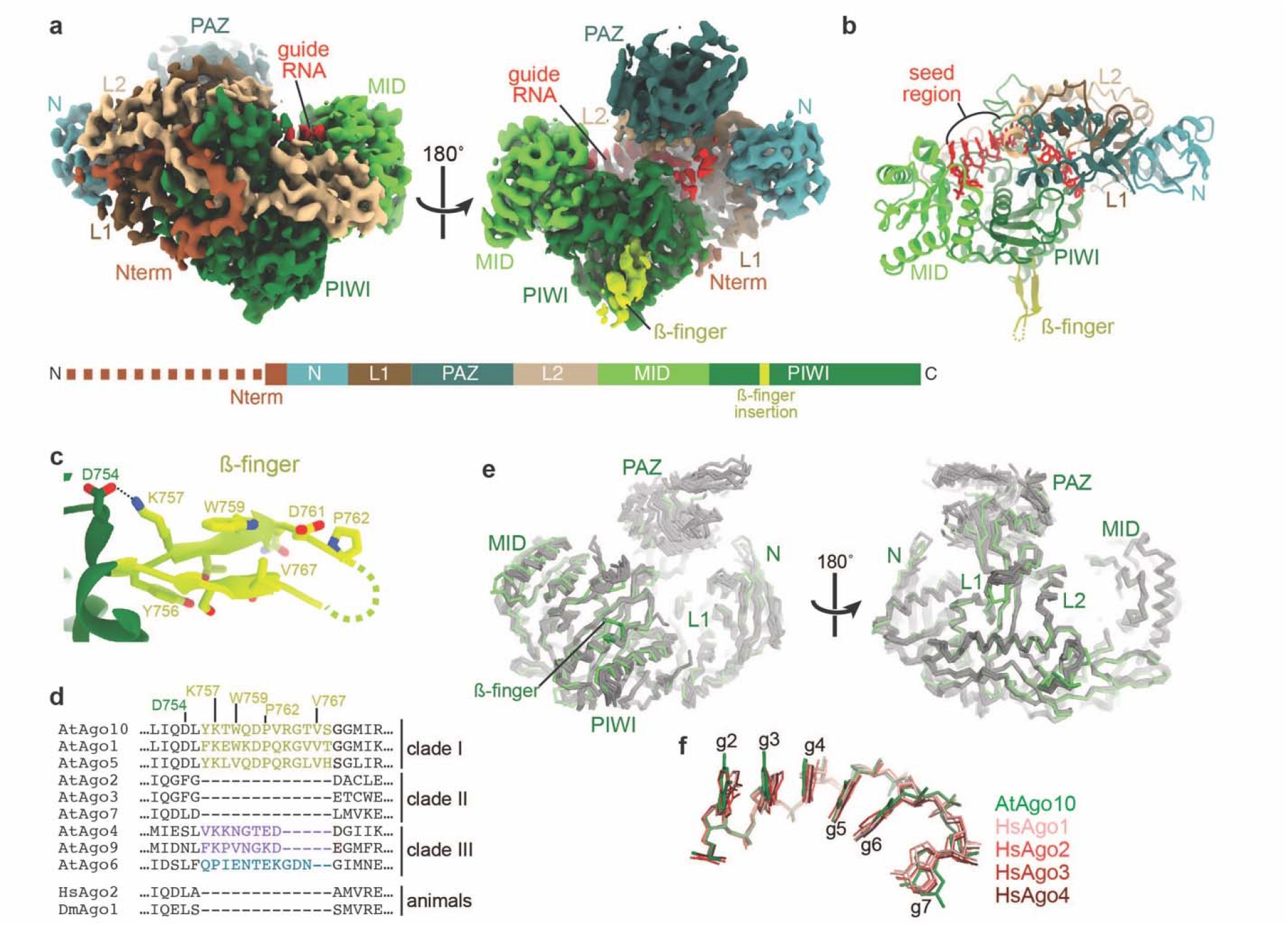
Structure of the AtAgo10-guide RNA complex. **a**. Cryo-EM reconstruction with individual domains segmented and colored as in schematic (lower panel). The dashed line indicates unstructured N-terminal residues. Guide RNA density colored red. **b**. Cartoon representation of the AtAgo10-guide RNA model. **c**. Close-up of the ß-finger structure. **d**. Sequence alignment near the ß-finger insertion of AGOs from *Arabidopsis*, human, and *Drosophila*. Plant AGO clades indicated. **e**. Superposition of AtAgo10-guide (green) and known human AGO-guide (silver) Ca backbone structures. **f**. Superposition of guide nucleotides (shown as sticks) from the seed regions of AtAgo10 (green) and representatives of the four human AGOs (shades of red).

### AtAgo10-guide-target complex adopts both slicing-competent and incompetent conformations

We next examined the ternary complex of AtAgo10 bound to a guide RNA and target with complementary to guide nucleotides g2-g16 by cryo-EM (Fig. S3). A catalytically inactive AtAgo10 mutant (D795A) was used to observe conformations that occur after target recognition but before target slicing. Initial 3D reconstructions indicated the presence of multiple RNA configurations within the AtAgo10 central cleft, leading to the refinement of two structures of the ternary AtAgo10-guide-target complex at ∼3.8 Å overall resolution (Fig. 2).

**Fig. 2.**
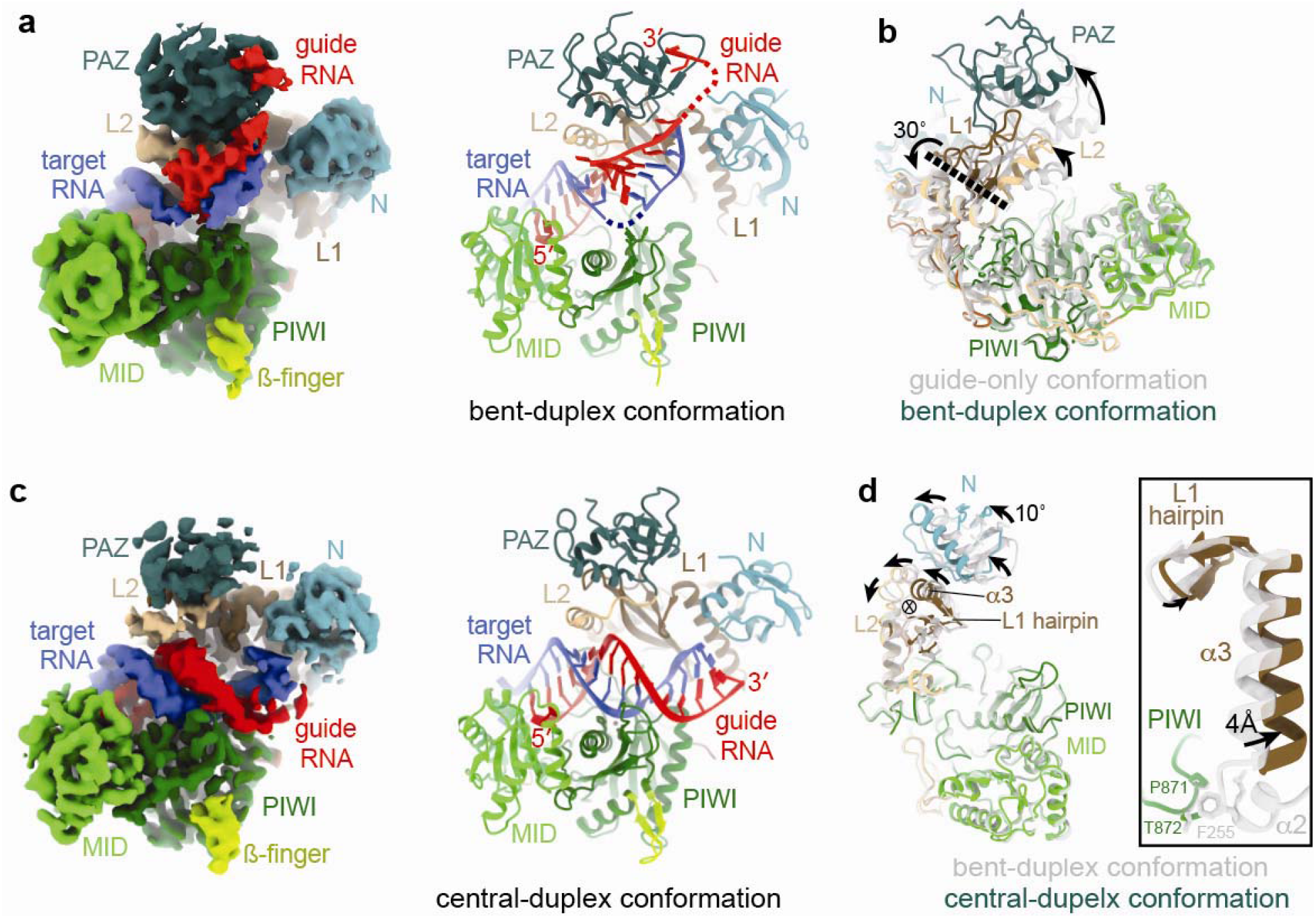
AtAgo10-guide-target structures. **a**. Reconstruction (left) and model (right) of the AtAgo10-guide-target complex in the bent-duplex conformation. **b**. Superposition of AtAgo10 guide-only (grey/translucent) and bent-duplex (colored) conformations (RNAs not shown). Arrows indicate movement directions from guide-only to bent-duplex structure. Dashed line indicates the axis of rotation (viewed from the side). **c**. Reconstruction (left) and model (right) of the AtAgo10-guide-target complex in the central-duplex conformation. **d**. Superposition of AtAgo10 bent-duplex (grey/translucent) and central-duplex (colored) conformations (RNAs and PAZ domain not shown). Arrows indicate movement directions from the bent-duplex to the central-duplex structure. Circled X indicates the axis of rotation (viewed down axis). Inset shows a closeup of the L1 hairpin, *α*3, and *α*2. *α*2 is disordered in the central-duplex structure, resulting in a loss of L1-PIWI contacts upon moving from the bent-duplex to the central-duplex conformation.

In the first structure (bent-duplex conformation), guide and target RNAs form a discontinuous duplex, with base pairing interrupted at guide nucleotides g9–g11 (Fig. 2a, S4). The unpaired segment forms a bend in the duplex that allows the guide 5’ and 3’ ends to remain anchored to nucleotide-binding pockets in the AtAgo10 MID and PAZ domains, respectively. Density for the guide-target duplex preceding the bend resolves its helical shape but is blurry, indicating the reconstruction represents a heterogenous family of guide-target interactions centered around pairing at g13–g15. Target-binding is associated with widening of the AtAgo10 central cleft via rotation of the PAZ domain about a hinge in the L1 and L2 domains (Fig. 2b). AtAgo10 conformation in the bent-duplex structure is nearly identical to a previous bent-duplex crystal structure of HsAgo2^20^, revealing conformational changes driven by extended guide-target pairing are conserved between plant and animal AGOs (Fig. S4f). Target (t) nucleotides t9-t11 (where t9 is the target nucleotide that is complementary to guide nucleotide g9) are disordered. Therefore, as slicing activity involves hydrolysis of the phosphodiester bond between t10 and t11^27^, the bent-duplex structure represents a family of related AtAgo10-guide-target conformations that are incapable of target slicing.

The second structure (central-duplex conformation) has density for a continuous guide-target RNA duplex passing through the AtAgo10 central cleft with the guide 3’ end released from the PAZ domain (Fig. 2c). The target RNA scissile phosphate is bound to catalytic residues D709 and H935 through a Mg^2+^ ion in a configuration resembling the catalytic centers of *T. thermophilus* AGO and bacterial RNase H^2,28^ (Fig. S5), indicating this reconstruction represents a slicing-competent conformation. AtAgo10 structure is similar to the bent-duplex conformation, with the exception of increased mobility in the PAZ domain (likely due to guide 3’ end release), and a ∼10° rotation of the N domain (Fig. 2d). The N domain rotates about an axis in the L1 domain near a conserved ß-hairpin structure (L1 hairpin), which also rotates along with portions of the L2 domain and helix-3 (*α*3) of the L1 domain. These conformational changes break contacts between the L1 and PIWI domains and are necessary to avoid steric clashes between the protein and target nucleotides after t13 (Fig. S6).

### Contacts to the target RNA in the slicing-competent conformation

The central-duplex structure reveals protein-RNA interactions associated with target slicing. AtAgo10 holds the guide-target duplex through contacts to both guide and target RNA backbones (Fig. 3a). In the seed region, the AtAgo10 MID, PIWI, and L2 domains contribute contacts to the guide backbone. These contacts are also present in the bent-duplex structure and similar to interactions in target-bound HsAgo2 crystal structures^19,20,29^, indicating a conserved role in small RNA targeting.

**Fig. 3.**
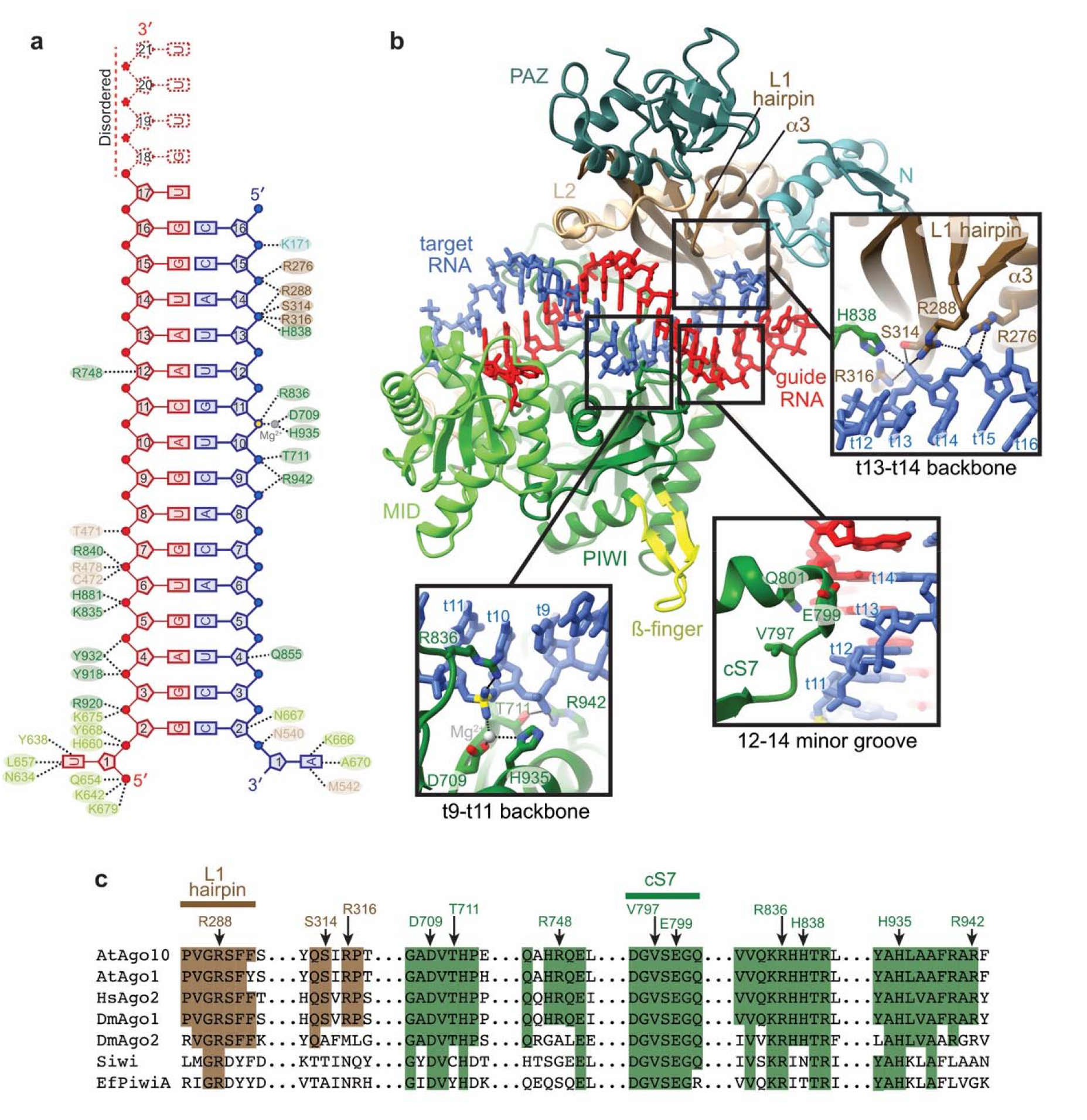
Catalytic conformation of the AtAgo10-guide-target complex. **a**. Schematic of major contacts between AtAgo10 and the guide (red) and target (blue) RNAs. Residues colored by domain, as in Fig. 1. **b**. Structure of the AtAgo10-guide-target central-duplex conformation. Insets detail protein-RNA contacts specific to the central-duplex conformation. **c**. Sequence alignment of select plant and animal AGO and PIWI proteins. AtAgo10 residues contacting t9-t14 are indicated. Residues forming L1 hairpin and cS7 are also indicated. Shaded residues are identical to AtAgo10.

Contacts unique to the central-duplex structure occur downstream of the seed region, where AtAgo10 binds the target RNA backbone using residues from the PIWI and L1 domains, including the L1 hairpin, which forms salt linkages with the phosphates of t13 and t14 (Fig. 3b). Additional contacts are made by a loop in the PIWI domain, previously termed ‘conserved sequence 7’ (cS7), which docks with the minor groove of the guide-target duplex at positions 12-14. AtAgo10 residues that recognize the backbone of target nucleotides t9-t14 are perfectly conserved in HsAgo2 (Fig. 3c).

### Docking of t9-t13 into the active site channel triggers target slicing

Based on the central-duplex structure, we propose a model for target recognition during RNA slicing. Previous studies showed mouse Ago2 slicing rates are most potently reduced by single or double mismatches in a region spanning t9–t13^16^. A surface representation of AtAgo10 shows the backbone of t9-t13 docks into a narrow channel containing the AtAgo10 active site (Fig. 4a). This ‘active site channel’ has shape complementarity to A-form RNA. Thus, we propose that docking of paired target nucleotides t9-t13 into the active site channel is the primary means by which Argonaute positions target RNA scissile phosphate for catalysis. The end of the active site channel is formed by cS7, explaining why cS7 is essential for RNA slicing^23^.

**Fig. 4.**
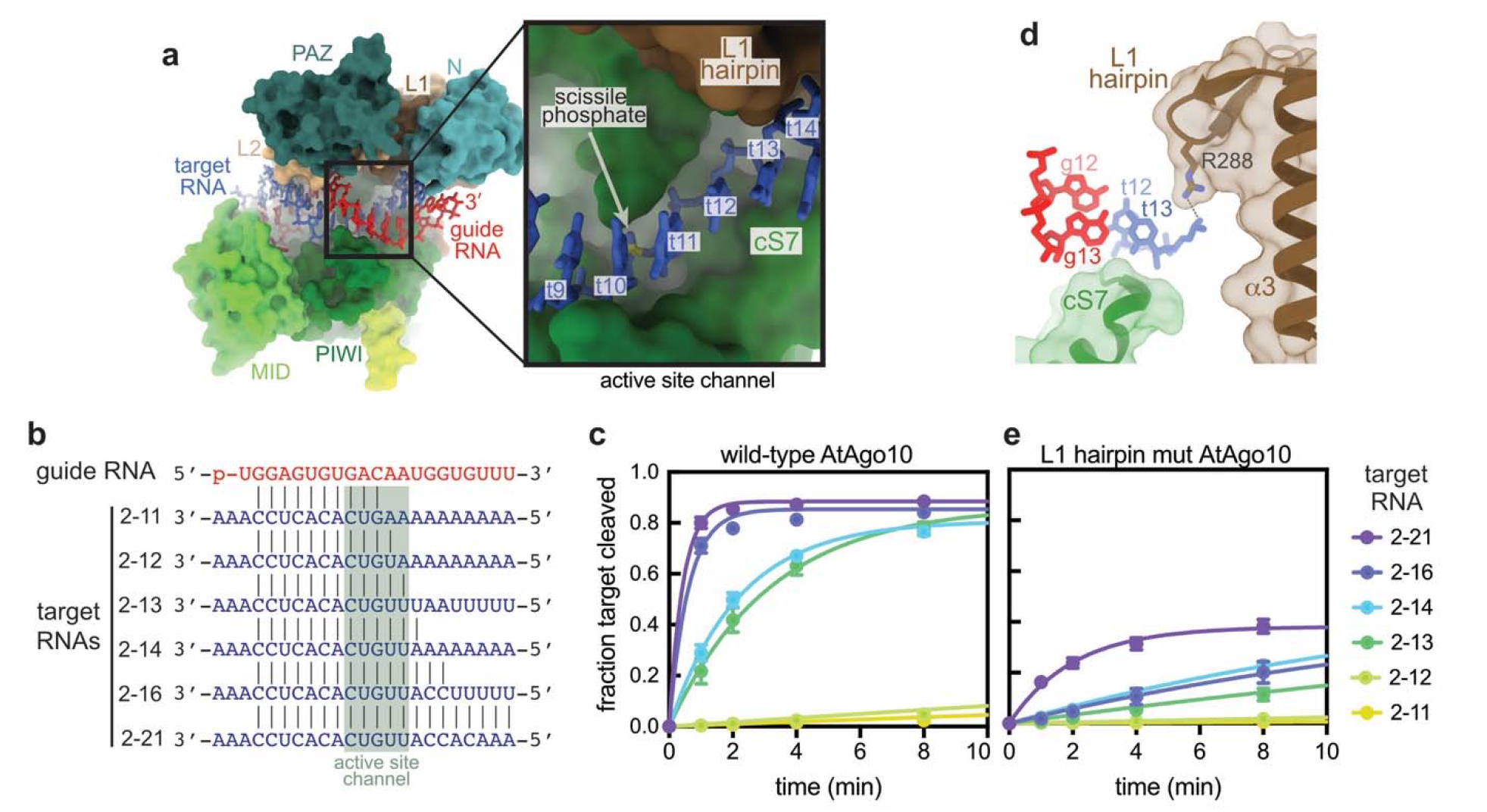
Structural basis for RNA Slicing. **a**. Surface representation of the AtAgo10 central-duplex model. Inset shows t9–t13 (blue sticks) docked in the active site channel (guide RNA omitted for clarity). **b**. Base pairing schematic of guide and target RNAs used in slicing reactions in panels c and e. Target nucleotides falling within the active site channel are indicated. **c**. Fraction of target RNAs cleaved by a saturating excess of wild-type AtAgo10-guide complex versus time. **d**. Side-view of the end of the active site channel where the L1 hairpin secures t13 against cS7. **e**. Fraction of target RNAs cleaved by a saturating excess L1-hairpin-mut AtAgo10-guide complex versus time. Data points are the mean values of three experiments. Error bars indicate SEM.

To test this model, we measured single turnover slicing rates by AtAgo10 using targets with increasing complementarity to the guide 3’ end (Fig. 4b). Slicing rates increased 40-fold as guide-target complementarity extended from t2-t12 to t2-t13, supporting the importance of pairing within the t9-t13 segment (Fig. 4c). Increasing from t2-t13 to t2-t14 had no effect on slicing rate, suggesting energy gains from t14 docking and pairing are used to stabilize the opened conformation of the L1/N domain (Fig. 2d, S6). Increasing complementarity to t2-t16 stimulated slicing rate an additional 5-fold, likely by stabilizing t9-t13 pairing and facilitating the release of the guide RNA 3’ end from the PAZ domain, which is necessary to dock the paired t9–t13 segment.

### The L1 hairpin braces targets in the active site channel

The central-duplex structure indicates that the L1 hairpin holds the t9-t13 segment in the active site channel (Fig. 4d). Indeed, an AtAgo10 mutant (L1-hairpin-mut AtAgo10), in which the apical loop of the L1 hairpin was replaced with a Gly-Gly linker, cleaved target RNAs ∼10 times slower than wild-type AtAgo10 (Fig 4e). Inhibition of slicing was not due to a target-binding defect as all target RNAs were bound by L1-hairpin-mut AtAgo10 (Fig. S7). Additionally, unlike wild-type AtAgo10, the t2-t14 and t2-t16 targets were cleaved at identical rates by L1-hairpin-mut AtAgo10. Taken with the conformational coupling between the L1 hairpin and N domain (Fig. 2d), we suggest the L1 hairpin coordinates docking of the paired t9–t14 segment with rotation of the N domain and thereby enables guide-target pairing downstream of position 14 in the slicing-competent conformation (Fig. S6). Finally, although >90% of the 2-21 target RNA molecules in our slicing reactions were bound to L1-hairpin-mut AtAgo10, <50% were cleaved after extended incubation (Fig. S7). Thus, by securing t9–t13 in the active site channel and coordinating target docking with downstream guide-target pairing, the L1 hairpin helps prevent AtAgo10-guide-target complexes from becoming trapped in stable, non-catalytic conformations.

### Structural basis for RNA slicing and functional implications

The model for slicing by TtAgo involves 5’-3’ propagation of the guide-target duplex driving extensive conformational changes in the relatively flexible TtAgo PIWI domain (Fig. S8a) to achieve a slicing-competent conformation^1,2^. By contrast, slicing by AtAgo10 involves a flexible guide-target duplex that is directed into the active site channel by large movements of relatively rigid domains in AtAgo10 (Fig. 2, S8b). We propose a ‘binder-clip’ mechanism for RNA slicing. In the guide-only conformation AGO is held closed, like a binder clip, with the L1 hairpin blocking the active site channel. Formation of the guide-target duplex pushes the AGO central cleft, including the L1 harpin, into an open conformation, like a binder clip when force is applied to open it. Due to the tension in the AGO protein, the L1 hairpin tends to collapse back towards the RNA, holding the guide-paired target strand in position for cleavage, like a binder clip holding paper once released. AGO structure (Fig. 1e,f), conformations (Fig. S4f), and RNA-contacting residues (Fig. 3c) are shared between AtAgo10 and HsAgo2, indicating this model may describe a conserved RNA slicing mechanism used in both Plantae and Animalia kingdoms of life.

The revised view of RNA slicing has implications for understanding siRNA potency. siRNA efficacy has long been known to be influenced by guide strand selection^30,31^ and target site accessibility^32^, and yet even after taking these factors into consideration highly potent siRNA sequences are relatively rare^33^. Our results indicate that the distribution of catalytic and non-catalytic conformations adopted by the AGO-guide-target complex is an additional factor that influences siRNA potency, wherein siRNA sequences that are prone to dwelling in bent-duplex conformations will be less potent than siRNAs that favor the central-duplex conformation. Thus, structural insight into AGO-guide-target conformations and the critical role of t9–t13 docking provides a rational basis for considering new motifs and modifications that dictate siRNA efficacy.

## Acknowledgments

We are grateful to Dr. Wenrong He and Dr. Yingnan Hou for plasmids containing *Arabidopsis thaliana* AGO cDNAs and to MacRae lab members, Guy Riddihough, and Jamie Williamson for advice. The research was funded by NIH grants R35GM127090 (I.J.M.), R01GM092740 (T.O.), and S10OD021634.

## Author contributions

Y.X. prepared AtAgo10 samples, performed biochemical experiments, produced high-resolution reconstructions, built AtAgo10 models, analyzed data, and co-wrote the manuscript. S.M. prepared cryo-EM grids and collected cryo-EM data. T.O. advised cryo-EM data collection and production of high-resolution reconstructions and provided mechanistic insights. I.J.M. provided structural and mechanistic insights, analyzed data, and co-wrote the manuscript.

## Data availability

Maps for the AtAgo10-guide, AtAgo10-guide-target (central-duplex), and AtAgo10-guide-target (bent-duplex) complexes were deposited in the Electron Microscopy Data Bank under accession IDs EMD-25446, EMD-25472, EMD-25482, respectively. Corresponding atomic models were deposited in the Protein Data Bank under accession IDs 7SVA, 7SWF, 7SWQ.

## References

1 Wang, Y. et al. Nucleation, propagation and cleavage of target RNAs in Ago silencing complexes. Nature 461, 754–761, doi:10.1038/nature08434 (2009).

2 Sheng, G. et al. Structure-based cleavage mechanism of Thermus thermophilus Argonaute DNA guide strand-mediated DNA target cleavage. Proceedings of the National Academy of Sciences of the United States of America 111, 652–657, doi:10.1073/pnas.1321032111 (2014).

3 Singh, A. et al. Plant small RNAs: advancement in the understanding of biogenesis and role in plant development. Planta 248, 545–558, doi:10.1007/s00425-018-2927-5 (2018).

4 Bartel, D. P. Metazoan MicroRNAs. Cell 173, 20–51, doi:10.1016/j.cell.2018.03.006 (2018).

5 Iwakawa, H. O. & Tomari, Y. Life of RISC: Formation, action, and degradation of RNA-induced silencing complex. Molecular cell 82, 30–43, doi:10.1016/j.molcel.2021.11.026 (2022).

6 Zamore, P. D., Tuschl, T., Sharp, P. A. & Bartel, D. P. RNAi: double-stranded RNA directs the ATP-dependent cleavage of mRNA at 21 to 23 nucleotide intervals. Cell 101, 25–33, doi:10.1016/S0092-8674(00)80620-0 (2000).

7 Hu, B. et al. Therapeutic siRNA: state of the art. Signal Transduct Target Ther 5, 101, doi:10.1038/s41392-020-0207-x (2020).

8 Llave, C., Xie, Z., Kasschau, K. D. & Carrington, J. C. Cleavage of Scarecrow-like mRNA targets directed by a class of Arabidopsis miRNA. Science 297, 2053–2056, doi:10.1126/science.1076311 (2002).

9 Allen, E., Xie, Z., Gustafson, A. M. & Carrington, J. C. microRNA-directed phasing during trans-acting siRNA biogenesis in plants. Cell 121, 207–221, doi:10.1016/j.cell.2005.04.004 (2005).

10 Axtell, M. J., Jan, C., Rajagopalan, R. & Bartel, D. P. A two-hit trigger for siRNA biogenesis in plants. Cell 127, 565–577, doi:10.1016/j.cell.2006.09.032 (2006).

11 Creasey, K. M. et al. miRNAs trigger widespread epigenetically activated siRNAs from transposons in Arabidopsis. Nature 508, 411–415, doi:10.1038/nature13069 (2014).

12 Ober-Reynolds, B. et al. High-throughput biochemical profiling reveals functional adaptation of a bacterial Argonaute. Molecular cell 82, 1329–1342 e1328, doi:10.1016/j.molcel.2022.02.026 (2022).

13 Wang, Y. et al. Structure of an argonaute silencing complex with a seed-containing guide DNA and target RNA duplex. Nature 456, 921–926, doi:10.1038/nature07666 (2008).

14 Wang, Y., Sheng, G., Juranek, S., Tuschl, T. & Patel, D. J. Structure of the guide-strand-containing argonaute silencing complex. Nature 456, 209–213, doi:10.1038/nature07315 (2008).

15 Wee, L. M., Flores-Jasso, C. F., Salomon, W. E. & Zamore, P. D. Argonaute divides its RNA guide into domains with distinct functions and RNA-binding properties. Cell 151, 1055–1067, doi:10.1016/j.cell.2012.10.036 (2012).

16 Becker, W. R. et al. High-Throughput Analysis Reveals Rules for Target RNA Binding and Cleavage by AGO2. Molecular cell, doi:10.1016/j.molcel.2019.06.012 (2019).

17 Niaz, S. The AGO proteins: an overview. Biol Chem 399, 525–547, doi:10.1515/hsz-2017-0329 (2018).

18 Pourjafar-Dehkordi, D. & Zacharias, M. Binding-induced functional-domain motions in the Argonaute characterized by adaptive advanced sampling. PLoS Comput Biol 17, e1009625, doi:10.1371/journal.pcbi.1009625 (2021).

19 Sheu-Gruttadauria, J., Xiao, Y., Gebert, L. F. & MacRae, I. J. Beyond the seed: structural basis for supplementary microRNA targeting by human Argonaute2. The EMBO journal, doi:10.15252/embj.2018101153 (2019).

20 Sheu-Gruttadauria, J. et al. Structural Basis for Target-Directed MicroRNA Degradation. Molecular cell, doi:10.1016/j.molcel.2019.06.019 (2019).

21 Nakanishi, K., Weinberg, D. E., Bartel, D. P. & Patel, D. J. Structure of yeast Argonaute with guide RNA. Nature 486, 368–374, doi:10.1038/nature11211 (2012).

22 Schirle, N. T. & MacRae, I. J. The crystal structure of human Argonaute2. Science 336, 1037–1040, doi:10.1126/science.1221551 (2012).

23 Nakanishi, K. et al. Eukaryote-specific insertion elements control human ARGONAUTE slicer activity. Cell Rep 3, 1893–1900, doi:10.1016/j.celrep.2013.06.010 (2013).

24 Park, M. S. et al. Human Argonaute3 has slicer activity. Nucleic acids research 45, 11867–11877, doi:10.1093/nar/gkx916 (2017).

25 Park, M. S. et al. Multidomain Convergence of Argonaute during RISC Assembly Correlates with the Formation of Internal Water Clusters. Molecular cell 75, 725–740 e726, doi:10.1016/j.molcel.2019.06.011 (2019).

26 Faehnle, C. R., Elkayam, E., Haase, A. D., Hannon, G. J. & Joshua-Tor, L. The making of a slicer: activation of human Argonaute-1. Cell Rep 3, 1901–1909, doi:10.1016/j.celrep.2013.05.033 (2013).

27 Elbashir, S. M., Lendeckel, W. & Tuschl, T. RNA interference is mediated by 21- and 22-nucleotide RNAs. Genes & development 15, 188–200 (2001).

28 Nowotny, M., Gaidamakov, S. A., Crouch, R. J. & Yang, W. Crystal structures of RNase H bound to an RNA/DNA hybrid: substrate specificity and metal-dependent catalysis. Cell 121, 1005–1016, doi:10.1016/j.cell.2005.04.024 (2005).

29 Schirle, N. T., Sheu-Gruttadauria, J. & MacRae, I. J. Structural basis for microRNA targeting. Science 346, 608–613, doi:10.1126/science.1258040 (2014).

30 Schwarz, D. S. et al. Asymmetry in the assembly of the RNAi enzyme complex. Cell 115, 199–208 (2003).

31 Khvorova, A., Reynolds, A. & Jayasena, S. D. Functional siRNAs and miRNAs exhibit strand bias. Cell 115, 209–216 (2003).

32 Ameres, S. L., Martinez, J. & Schroeder, R. Molecular basis for target RNA recognition and cleavage by human RISC. Cell 130, 101–112, doi:10.1016/j.cell.2007.04.037 (2007).

33 Reynolds, A. et al. Rational siRNA design for RNA interference. Nature biotechnology 22, 326–330, doi:10.1038/nbt936 (2004).

