## Supplementary Materials for "Structural basis for RNA slicing by a plant Argonaute"

**Contents:**

Materials and Methods
Fig. S1. Imaging and processing of the AtAgo10-guide RNA complex.
Fig. S2. Details of AtAgo10 structural analysis
Fig. S3. Imaging and processing of the AtAgo10-guide-target RNA complex
Fig. S4. Reconstruction of the AtAgo10-guide-target bent-duplex conformation
Fig. S5. Reconstruction of the AtAgo10-guide-target central-duplex conformation
Fig. S6. Bent-duplex conformation of AtAgo10 cannot bind a central guide-target duplex with pairing beyond t13
Fig. S7. L1-hairpin-mut AtAgo10 is impaired in target-slicing but not target-binding
Fig. S8. The changes in TtAgo and AtAgo10 PIWI domain by forming slicing competent conformation
Table S1. Cryo-EM data collection, refinement, and validation statistics

### Methods and Materials

Bacterial strains and plasmids

Bacteria used for cloning were chemically competent *E. coli* OmniMAX$^{TM}$ (Thermo Fisher). Bacteria used for production of bacmid were DH10Bac$^{TM}$ chemically competent *E. coli* (Thermo Fisher).

Bacterial media and growth conditions

All bacterial cultures were grown in Luria-Bertani (LB) medium at 37 °C. When needed, media was supplemented with one or more of the following antibiotics at the following concentrations: ampicillin (100 µg/mL), kanamycin (40 µg/mL), tetracycline (5 µg/mL), gentamycin (7 µg/mL), 5-Bromo-4-Chloro-3-Indolyl β-D-Galactopyranoside (X-gal, 25 µg/mL in dimethylformamide), and/or Isopropyl β-D-1-thiogalactopyranoside (IPTG, 1 mM).

Insect cell media and growth conditions

Sf9 cells were grown in Lonza Insect XPRESS$^{TM}$ medium supplemented with 1x Gibco™ Antibiotic-Antimycotic in suspension at 27 °C.

Cloning and mutagenesis

DNA fragments encoding full length of AtAgo10 were generated by PCR using a cDNA clone encoding *Arabidopsis thaliana* ARGONAUTE 10 (TAIR: AT5G43810, a gift from Dr. Wenrong He and Dr. Yingnan Hou) as template. PCR products were cloned as SfoI-KpnI fragments into a modified form of pFastBac HTA (Thermo Fisher) to generate expression plasmids for the Bac-to-Bac baculovirus expression system (Thermo Fisher). The active site mutation D795A was generated by quick-change PCR. The L1 harpin mutation was generated by first amplifying two AtAgo10 fragments with PCR primers overlapping and replacing codons for the L1 harpin residues 286-289 (VGRS) with GG codons, and then assembling the two gel purified PCR fragments with digested pFastBac HTA by NEBuilder® HiFi DNA Assembly Master Mix.

Preparation of AtAgo10-guide RNA complex

$His_6$-Flag-Tev-tagged AtAgo10 proteins were expressed in Sf9 cells using a baculovirus system (Invitrogen). 60 hour cultured 750 mL 3.4 x $10^6$ cells/mL Sf9 cells infected with ~15 mL virus were harvested by centrifugation. Usually, two of these cell pellets were combined and resuspended in ~200 mL Lysis Buffer (50 mM NaH2PO4, pH 8, 300 mM NaCl, 5% glycerol, 20 mM imidazole, 0.5 mM TCEP). Resuspended cells were lysed by passing twice through a M-110P lab homogenizer (Microfluidics). The resulting total cell lysate was clarified by centrifugation (30,000 x g for 25 min) and the supernatant fraction was applied to 8 mL packed Ni-NTA resin (Qiagen) and gently rocked at 4˚C for 1.5 hours in 50 mL conical tubes. Resin was pelleted by brief centrifugation and the supernatant solution was discarded. The resin was washed with ~50 mL ice cold Nickel Wash Buffer (300 mM NaCl, 20 mM imidazole, 0.5 mM TCEP, 50 mM Tris, pH 8). Centrifugation/wash steps were repeated a total of three times. Co-purified cellular RNAs were degraded by incubating with 400U micrococcal nuclease (Clontech) on-resin in ~25 mL of Nickel Wash Buffer supplemented with 5 mM $CaCl_2$ at room temperature for ~1 hour. The nuclease-treated resin was washed three times again with Nickel Wash Buffer and then eluted in four column volumes of Nickel Elution Buffer (300 mM NaCl, 300 mM imidazole, 0.5 mM TCEP, 50 mM Tris, pH 8). Eluted protein was incubated with 24 nmol synthetic guide RNA and 150 µg TEV protease during a ~1 hour dialysis against 1 L of Dialysis Buffer (300 mM NaCl, 0.5 mM TCEP, 50 mM Tris, pH 8) at room temperature. During dialysis, a capture resin for isolation of guide-loaded AtAgo10 was prepared by incubating 345.6 uL packed High Capacity Neutravidin Resin (Thermo Fisher) with 28.8 nmol capture oligo in wash A buffer (100 mM KOAc, 2 mM MgOAc, 0.01% CHAPS, 30 mM Tris, pH 8) for 30 min at 4 ˚C, followed by 10 mL wash A buffer wash. The dialyzed RNA-protein sample was then supplemented with 0.01% CHAPS and 2 mM MgOAc and then incubated with capture resin at RT for 1 hour (without rocking!!!! Just gently inverting the tube every 5-10 minutes). The resin was then washed, in order, three times with 10 mL Wash A, four times with 10 mL Wash B (2 M KOAc, 2 mM MgOAc, 0.01% CHAPS, 30 mM Tris, pH 8), and three times with 10 mL Wash C (1 M KOAc, 2 mM MgOAc, 0.01% CHAPS, 30 mM Tris, pH 8). The resin was then re-suspended in 1900 µL Wash C supplemented with 57.6 nmol competitor DNA at RT for ~2 hours (Gently inverting the tube every 5-10 minutes). The eluate was recovered and dialyzed against 1 L Q dialysis buffer (150 mM NaCl, 0.01% CHAPS, 0.5 mM TCEP, 20 mM Tris, pH 8) at 4 ˚C overnight. After dialysis, the sample was passed through 240 µl of Q Sepharose Fast Flow anion exchange resin (GE Healthcare) (pre-equilibrated in Q dialysis buffer) to remove unbound oligonucleotides, and flow-through solution was collected. The flow-through was then concentrated to 1~3 mg/mL while buffer exchanging to Tris Crystal buffer (10 mM Tris pH 8, 100 mM NaCl, 0.5 mM TCEP). The concentrated protein was aliquoted, flash-frozen in liquid $N_2$ and stored at -80 ˚C. Concentration of the AtAgo10-guide RNA complex was determined by Bradford assay using BSA as a standard.

Grid preparation for cryo-EM

For AtAgo10-guide RNA binary complex grids, 3-4 µL of 0.3 mg/mL AtAgo10-guide RNA in Tris Crystal Buffer was added onto freshly plasma cleaned (75% nitrogen, 25% oxygen atmosphere at 15 W for 7 seconds in Solarus plasma cleaner, Gatan) 300 mesh holey gold grids (UltrAuFoil R1.2/1.3, Quantifoil). Excess sample solution was removed from grids by manual blotting with Whatman No.1 filter paper for 1-2 seconds. Samples were immediately vitrified by plunge freezing in liquid-ethane at -179 °C using a manual plunge freezing device. Grid vitrification was performed in a cold room maintained at 4°C with relative humidity between 90-98% to minimize sample evaporation. For AtAgo10-guide-target RNA ternary complex grids, 1.2 molar equivalents of target RNA was added to purified AtAgo10-miRNA complex, which was incubated on ice for ~1h in Tris Crystal Buffer with 2 mM $MgCl_2$. Grid preparation procedure was the same as the AtAgo10-guide RNA binary complex.

Cryo-EM data acquisition

Cryo-EM data acquisition was performed on a 200kV Talos Arctica (Thermo Fisher Scientific) transmission electron microscope. Micrographs were acquired using a K2 Summit (Gatan) direct electron detector, operated in electron-counting mode, using the automated data collection software Leginon (*32*) by image shift-based movements from the center of four adjacent holes to target the center of each hole for exposures. Each micrograph for the AtAgo10-guide RNA binary complex was collected as 55 dose-fractionated movie frames over 11 s and with a cumulative electron exposure of 66.51 e-/Å$^2$. The data set was collected at a nominal magnification of 73kx, corresponding to 0.566 Å/pixel on the detector, with random nominal defocus values varying between 0.8 µm and 1.3 µm. For the AtAgo10-guide-target RNA ternary complex, each micrograph was acquired as 42 dose-fractionated movie frames over 8.4 s with a cumulative electron exposure of 66.88 e-/Å$^2$. The data set was collected at a nominal magnification of 45kx, corresponding to 0.91 Å/pixel on the detector, with random nominal defocus values varying between 1 µm and 1.7 µm. 2,656 and 2,049 micrographs were collected for the AtAgo10-guide and AtAgo10-guide-target complexes, respectively.

Image processing and 3D reconstruction

For the Atago10-guide-target ternary complex map, raw movies were imported into the RELION 3.1 data processing pipeline(*33*). Beam-induced motion correction and radiation damage compensation over spatial frequencies (dose-weighting) of the raw movies, was performed using RELION embedded MotionCor2 with its own implementation. Contrast Transfer Function (CTF) parameters for these micrographs were estimated using gCTF. The micrographs with estimated max resolution lower than 7Å were eliminated for following processing. Laplacian of Gaussian based automated particle picking program in RELION was used for picking particles from the first 125 micrographs yielding a stack of 358,840 picks that were binned 4 x 4 (3.64 Å/pixel, 50-pixel box size) and subjected to reference-free 2D classification. The best four classes that represented orthogonal views of AtAgo10 were then used for template-based particle picking.1,935,081 picks were extracted from 1318 micrographs, binned 2 x 2 (1.82 Å/pixel, 100-pixel box size) and subjected to reference-free 2D classification using a 120 Å soft circular mask. The best 2D class averages that represented side or top/bottom views of AtAgo10 (i.e., the longest dimensions of AtAgo10) were then isolated (972,516 particles). Those class averages that contained “end-on” pr tilted views of AtAgo10 were combined and subjected to another round of reference-free 2D classification using a 80 Å soft circular mask to focus on alignment of the smaller “end-on” views. The best 2D class averages were then selected (75,722 particles) and combined with previously selected particles containing the longer side views for further processing. A total of 1,048,238 particles corresponding to the best 2D class averages that displayed strong secondary-structural elements and multiple views of AtAgo10 were selected for homogenous ab initio model generation using cryoSPARC (*34*). The generated model was imported back into RELION and low-pass filtered to 35 Å for use as an initial model for 3D classification with alignment (6 classes, tau_fudge = 4). For the bent duplex conformation (conformation-2), particles comprising the best-resolved class (Class 4) was then subjected to 3D auto-refinement. A subsequent no-alignment classification (2 classes, tau_fudge = 20) was performed and the class that possessed the best-resolved side-chain and backbone densities were re-centered and re-extracted unbinned (0.91 Å/pixel, 200-pixel box size). Due to the close proximity of neighboring particles, any re-centered particle within 40 Å of another was considered a duplicate and subsequently removed. This final stack of 28,499 particles was 3D auto-refined using a soft mask (0-pixel extension, 5-pixel soft cosine edge). Following per-particle defocus, beam-tilt refinement and Bayesian polishing, the final resolution of the reconstruction improved to 3.79 Å (gold-standard FSC at 0.143 cutoff) for this conformation. For central duplex conformation, particles comprising Class 4 and Class 6 was combined and subjected to another round of 3D classification with alignment (4 classes, tau_fudge = 4). Particles comprising the best-resolved class was then subjected to 3D auto-refinement. A subsequent no-alignment classification (2 classes, tau_fudge = 20) was performed and the class that possessed the best-resolved side-chain and backbone densities were re-centered and re-extracted unbinned (0.91 Å/pixel, 200-pixel box size). Due to the close proximity of neighboring particles, any re-centered particle within 40 Å of another was considered a duplicate and subsequently removed. The final stack of 38,833 particles was 3D auto-refined using a soft mask (0-pixel extension, 5-pixel soft cosine edge). Following per-particle defocus, beam-tilt refinement and Bayesian polishing, the final resolution of the reconstruction improved to 3.79 Å (gold-standard FSC at 0.143 cutoff) for this conformation.

For the AtAgo10-guide RNA binary complex dataset, beam-induced motion correction and radiation damage compensation over spatial frequencies (dose-weighting) of the raw movies, was performed using UCSF MotionCor2 (*35*) implemented in the Appion (*36*) image processing workflow. Motion corrected, summed micrographs were imported into the RELION 3.1 data processing pipeline. Contrast Transfer Function (CTF) parameters for these micrographs were estimated using gCTF. Laplacian of Gaussian based automated particle picking program in RELION was used for picking particles from random 843 micrographs yielding a stack of 716,944 picks that were binned 4 x 4 (2.264 Å/pixel, 72-pixel box size) and subjected to reference-free 2D classification. The best four classes that represented orthogonal views of AtAgo10 were then used for template-based particle picking.763,209 picks were extracted from 2645 micrographs, binned 2 x 2 (1.132 Å/pixel, 164-pixel box size) and subjected to reference-free 2D classification using a 120 Å soft circular mask. A total of 381,087 particles corresponding to the best 2D class averages that displayed strong secondary-structural elements and multiple views of AtAgo10 were isolated. Previous AtAgo10-guide-target RNA ternary complex map (bent duplex conformation) was down-scaled and low-pass filtered to 30 Å for use as an initial model for 3D auto-refinement. 381,087 particles were 3D auto-refined into a single class. Upon convergence, the run was continued with a soft mask (3-pixel extension, 5-pixel soft cosine edge), followed by subsequent re-centering and re-extraction binned 2 × 2 (1.132 Å/pixel, 164-pixel box size). Due to the close proximity of neighboring particles, any re-centered particle within a 40 Å of another was considered a duplicate and subsequently removed. These particles were then 3D auto-refinement into a single class using the same soft mask, followed by A subsequent no-alignment classification (2 classes, tau_fudge = 20) was performed and the class that possessed the best-resolved side-chain and backbone densities were re-centered and re-extracted unbinned (0.566 Å/pixel, 328-pixel box size). This final stack of 53,766 particles was 3D auto-refined using a soft mask (0-pixel extension, 5-pixel soft cosine edge). Following per-particle defocus and beam tilt refinement, the final resolution of the reconstruction improved to 3.26 Å (gold-standard FSC at 0.143 cutoff).

Model building and refinement

The homology model of AtAgo10 was generated using Human Ago2 structure (PDB: 4OLA) by Modeller(homology) in UCSF Chimera (*37, 38*). The initial homology model was then docked into the AtAgo10-guide RNA reconstruction using UCSF Chimera, followed by manual model building using Coot (*39*). The two AtAgo10-guide-target models were built in a similar fashion using the AtAgo10-guide RNA structure as an initial model. Models were refined through iterative rounds of manual building and fixing of geometric and rotameric outliers in Coot and real-space refinement optimizing global minimization, atomic displacement parameters and local grid search using PHENIX (*40*). Model validation was performed using MolProbity (http://molprobity.biochem.duke.edu) and PDB validation servers (https://www.rcsb.org). Structural figures were made using PyMOL (Schrödinger, LLC) and UCSF ChimeraX.

Target Slicing assay

Purified AtAgo10-guide and AtAgo10-L1-mut-guide RNA complexes (10 nM, final concentration) were incubated at room template with complementary $^{32}$P 5'-radiolabelled target RNAs (2 nM, final concentrations) in reaction buffer composed of 30 mM Tris pH 8.0, 100 mM potassium acetate, 2 mM Magnesium acetate, 0.5 mM TCEP, and 0.01 mg/mL baker's yeast tRNA. Target slicing reactions were stopped at various times by mixing aliquots of each reaction with an equal volume of denaturing gel loading buffer (98% w/v formamide, 0.025% xylene cyanol, 0.025% w/v bromophenol blue, 10 mM EDTA pH 8.0). Intact and cleaved target RNAs were resolved by denaturing PAGE (15%) and visualized by phosphorimaging. Quantification of $^{32}$P signal was performed using ImageQuant TL (GE Healthcare). Fraction target RNA cleaved was calculated by dividing the $^{32}$P signal in bands on the gel corresponding to the cleavage products by the sum of the $^{32}$P signals in intact and cleavage bands for each target RNA at each time point.

Target binding assays

Purified AtAgo10-L1-mut-guide RNA complexes (10 nM, final concentration) were incubated at room template for 1 hour with complementary $^{32}$P 5'-radiolabelled target RNAs (2 nM, final concentrations) in a reaction buffer composed of 30 mM Tris pH 8.0, 100 mM potassium acetate, 0.5 mM TCEP, 0.01 mg/mL baker's yeast tRNA and 0.005% NP-40. $Mg^{2+}$ was excluded from the reaction buffer to prevent target slicing. Using a dot-blot apparatus (GE Healthcare Life Sciences), protein-RNA complexes were captured on Protran nitrocellulose membrane (0.45 µm pore size, Whatman, GE Healthcare Life Sciences) and unbound RNA on Hybond Nylon membrane (Amersham, GE Healthcare Life Sciences). Samples were applied with vacuum and immediately washed once with 100 µL of ice-cold wash buffer (30 mM Tris pH 8.0, 100 mM potassium acetate, 0.5 mM TCEP). Membranes were air-dried and $^{32}$P signal was visualized and by phosphorimaging and quantified using ImageQuant TL (GE Healthcare). Fraction target RNA bound was calculated by dividing the $^{32}P$ signal retained on the nitrocellulose membrane by the sum of the $^{32}P$ signals on nitrocellulose and nylon membranes for each target RNA.

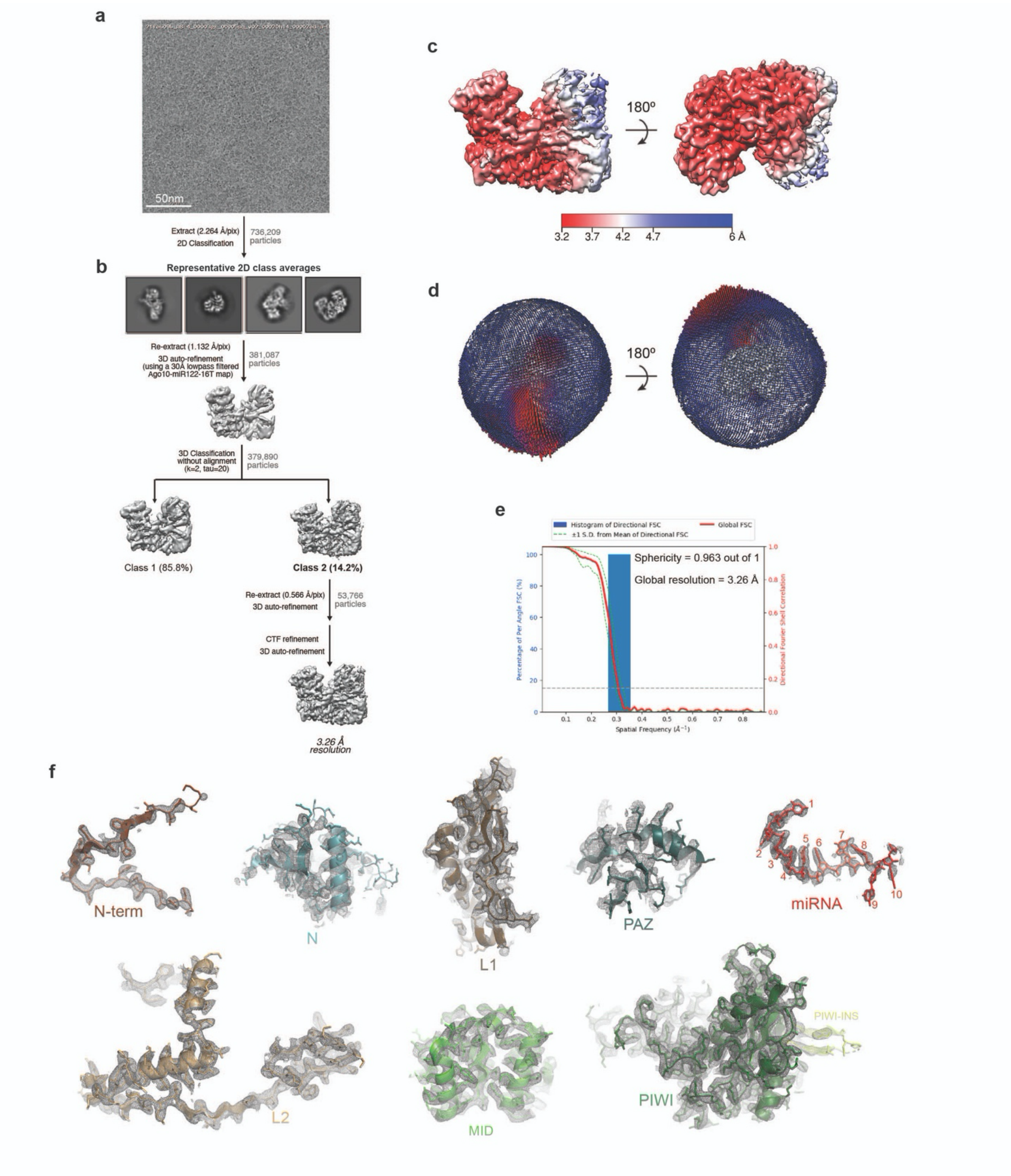

**Fig. S1. Imaging and processing of the AtAgo10-guide RNA complex.**
**a.** Representative cryo-EM micrograph. **b.** Cryo-EM data processing workflow. Particles isolated from micrographs were sorted by reference-free 2D classification. Only particles containing high-resolution features for the intact complex were selected for downstream processing. 3D classification was used to further remove low-resolution or damaged particles, and the remaining particles were refined to obtain a 3.26 Å resolution reconstruction. **c.** The final 3D map for the AtAgo10-guide RNA complex colored by local resolution values, where the majority of the map was resolved between 3.3 Å and 3.7 Å and the map covering the more mobile PAZ and N domains

having lower resolution. **d.** Angular distribution plot showing the Euler angle distribution of the AtAgo10-guide particles in the final reconstruction. The position of each cylinder corresponds to the 3D angular assignments and their height and color (blue to red) corresponds to the number of particles in that angular orientation. **e.** Directional Fourier Shell Correlation (FSC) plot representing 3D resolution anisotropy in the reconstructed map, with the red line showing the global FSC, green dashed lines correspond to ±1 standard deviation from mean of directional resolutions, and the blue histograms correspond to percentage of directional resolution over the 3D FSC. **f.** EM density quality of AtAgo10-guide RNA complex. Individual domains of AtAgo10 fit into the EM density, EM density shown in mesh; molecular models (colored as in Fig. 1) shown in cartoon representation with side chains shown as sticks; guide RNA shown in stick representation.

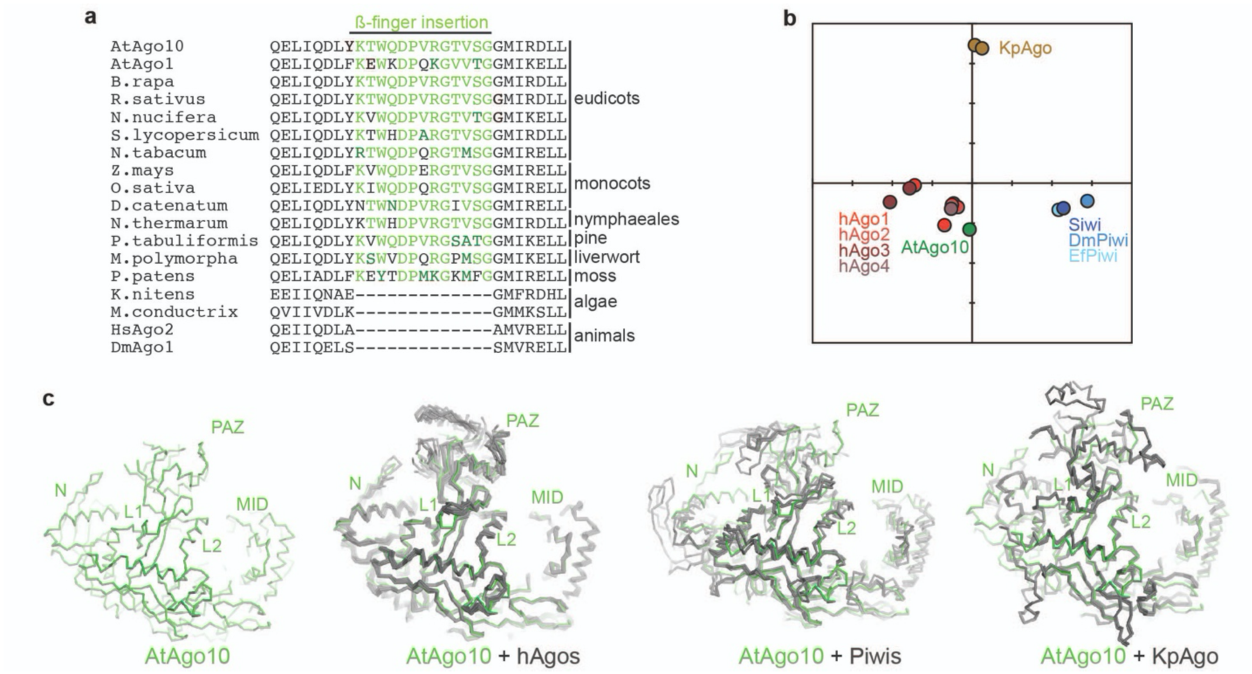

**Fig. S2. Details of AtAgo10 structural analysis**

**a.** Alignment of ß-finger sequences from clade I AGOs from diverse plants, algae, and animals. The insertion can be found in clade I AGOs in all plant phyla but is not apparent in animals or algae, indicating β-finger arose around the emergence of land plants and has been broadly maintained in the clade I AGOs over the last ~500 million years. **b.** All-against-all correspondence analysis (*62*) of known eukaryotic AGO and PIWI guide-bound structures (target-bound structures not included). **c.** C$\alpha$ alignment of AtAgo10 with known human AGO (PDB: 4KRF, 4KXT, 4OLA, 4W5N, 5JS1, 5VM9, 6OON), PIWI (PDB: 5GUH, 6KR6, 7KX7), and yeast AGO (PDB: 4F1N) guide-bound structures.

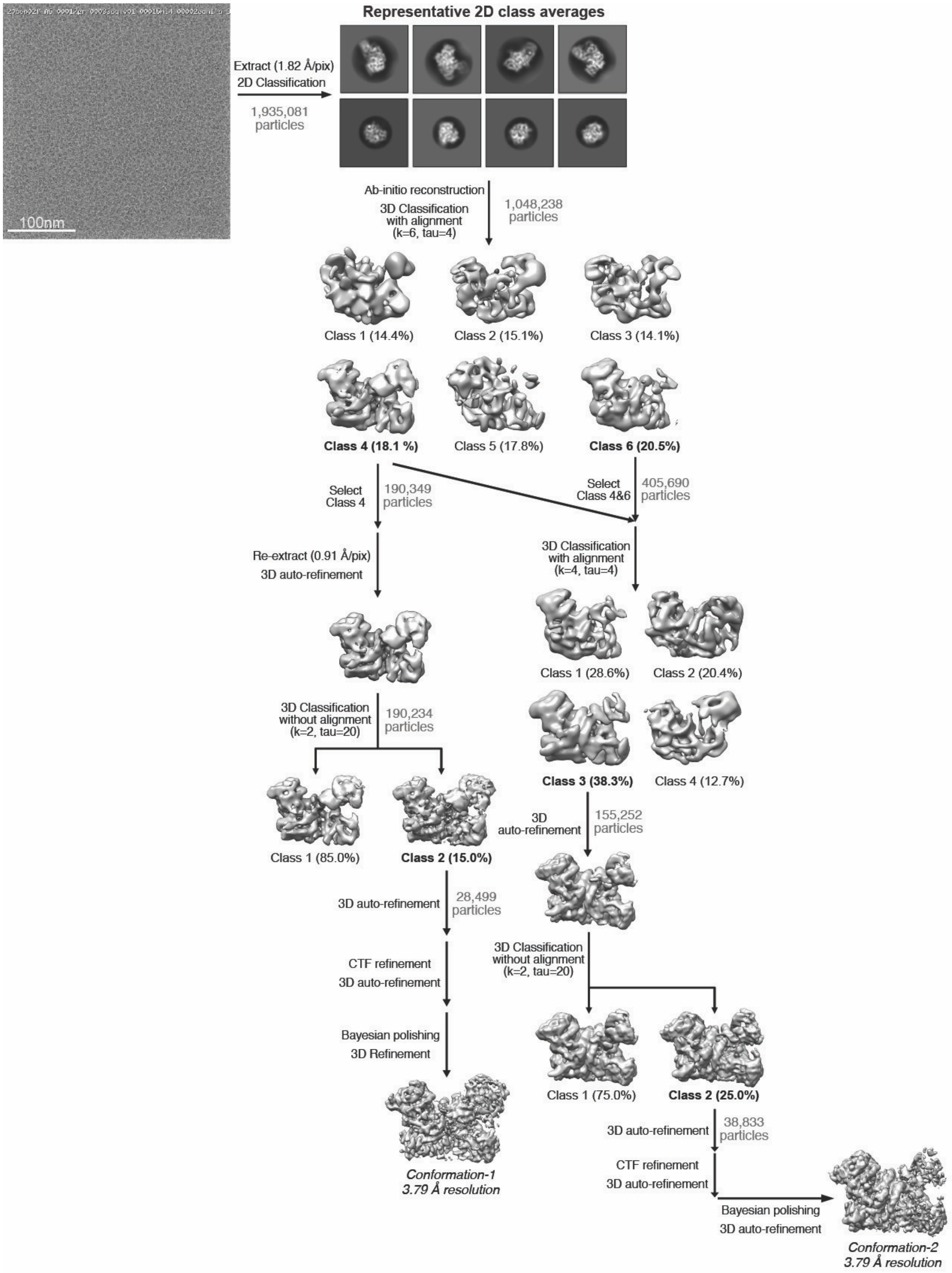

**Fig. S3. Imaging and processing of the AtAgo10-guide-target RNA complex**
Representative cryo-EM micrograph in upper left, proceeded by cryo-EM data processing workflow. Particles isolated from micrographs were sorted by reference-free 2D classification. Only particles containing high-resolution features for the intact complex were selected for downstream processing. 3D classification was used to further remove low-resolution or damaged particles, and isolate classes with distinct RNA conformations. 3D classification and particles in

the two major classes were further refined to obtain two 3.79 Å resolution reconstructions. Conformation-1 corresponds to the ‘bent-duplex’. Conformation-1 is the ‘central-duplex’, catalytic conformation.

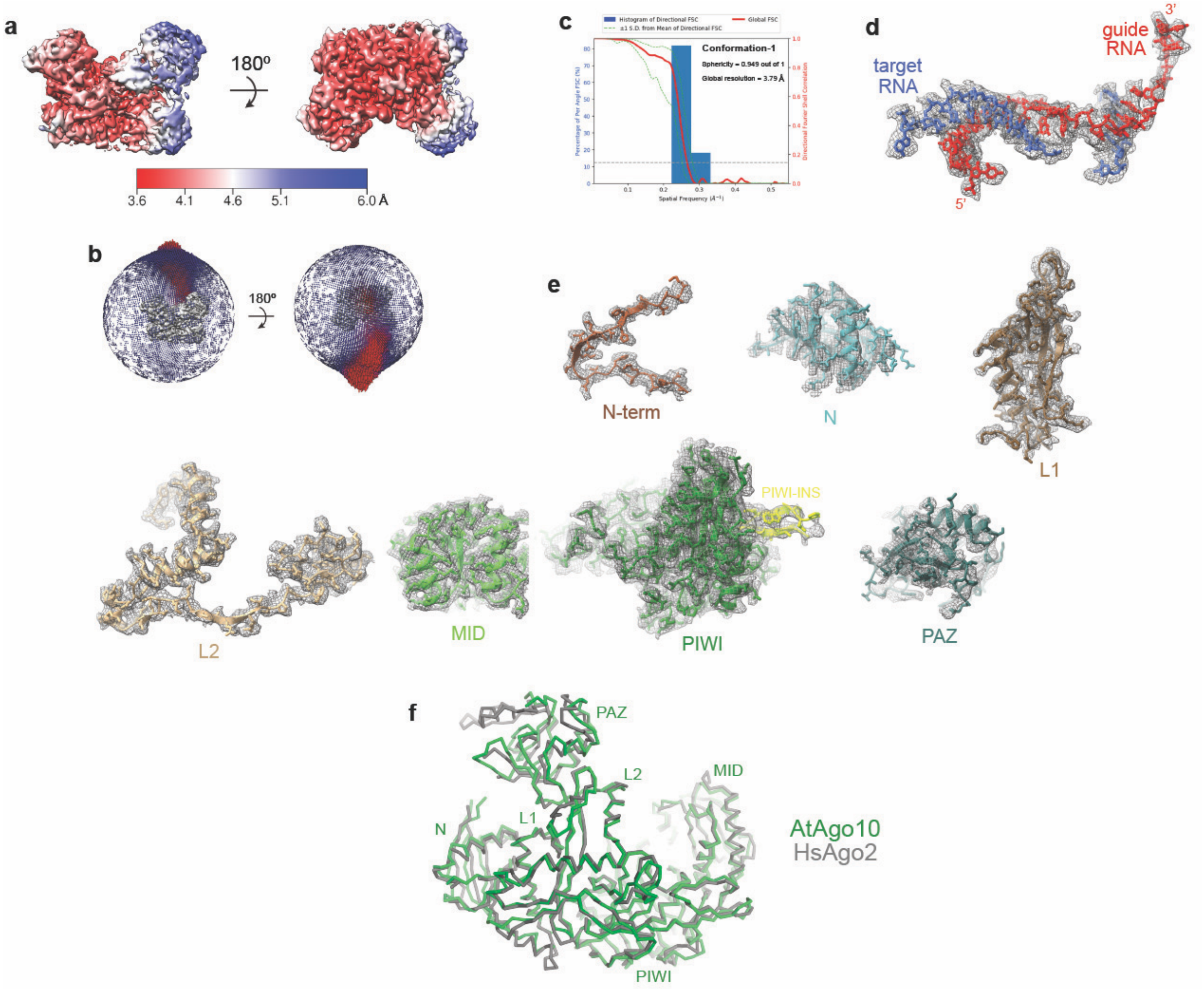

**Fig. S4. Reconstruction of the AtAgo10-guide-target bent-duplex conformation**
**a.** Final 3D map for Conformation-2 (see Fig. S4) of the AtAgo10-guide-target RNA complex colored by local resolution values. **b.** Angular distribution plot showing the Euler angle distribution of the AtAgo10-guide-target particles in the final reconstruction. The position of each cylinder corresponds to the 3D angular assignments and their height and color (blue to red) corresponds to the number of particles in that angular orientation. **c.** Directional Fourier Shell Correlation (FSC) plot representing 3D resolution anisotropy in the reconstructed map, with the red line showing the global FSC, green dashed lines correspond to ±1 standard deviation from mean of directional resolutions, and the blue histograms correspond to percentage of directional resolution over the 3D FSC. **d.** EM density quality of the guide-target duplex. **e.** Individual AtAgo10 domains fit into the EM density, EM density shown in mesh; molecular models (colored as in Fig. 1) shown in cartoon representation with side chains shown as sticks. **f.** The bent-duplex conformation of AtAgo10 strongly resembles a previous bent duplex structure of HsAgo2 (PDB 6MDZ), with an RMSD of 1.5 Å for 634 equivalent Cα atoms.

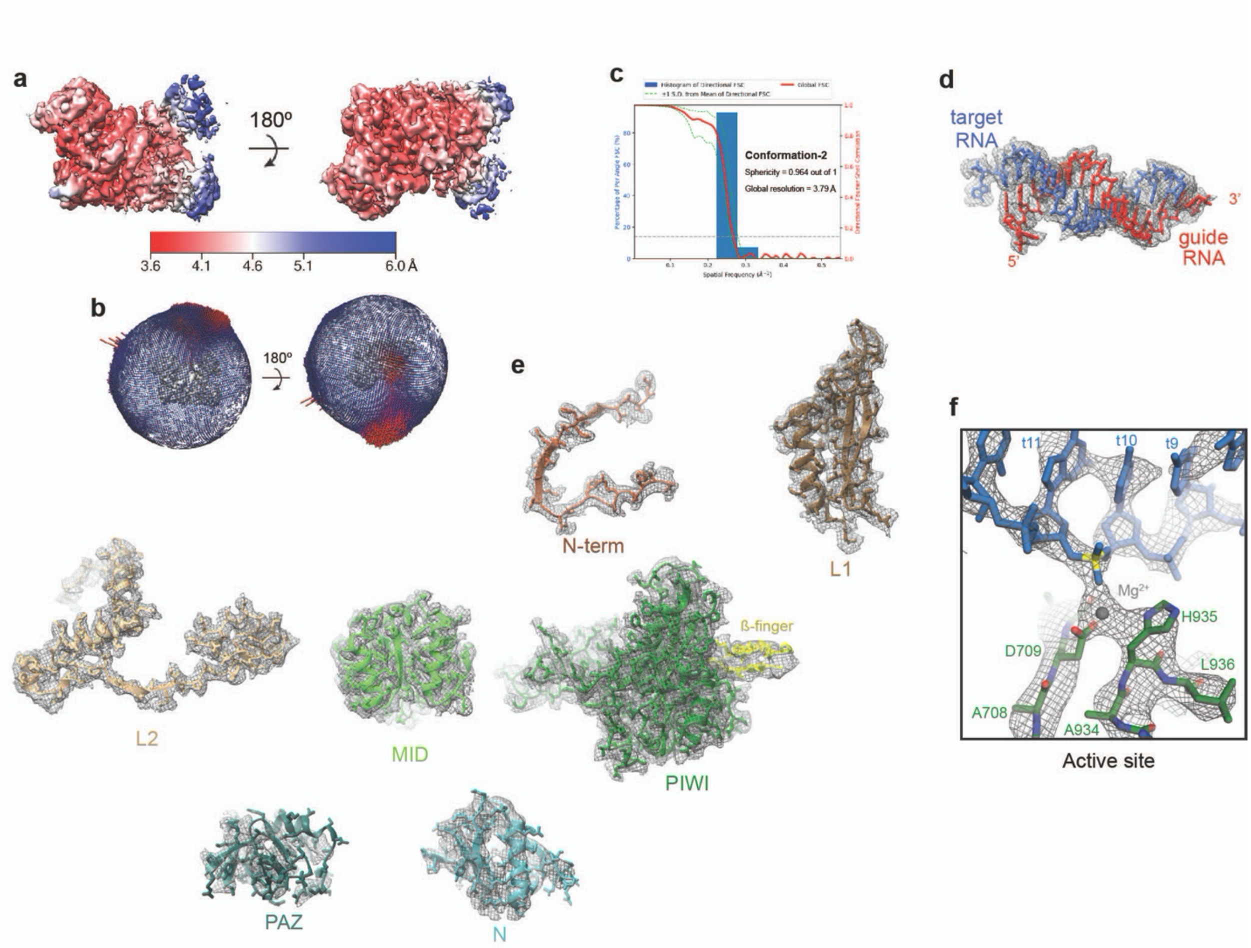

**Fig. S5. Reconstruction of the AtAgo10-guide-target central-duplex conformation**
**a.** Final 3D map for Conformation-1 (see Fig. S4) of the AtAgo10-guide-target RNA complex colored by local resolution values. **b.** Angular distribution plot showing the Euler angle distribution of the AtAgo10-guide-target particles in the final reconstruction. The position of each cylinder corresponds to the 3D angular assignments and their height and color (blue to red) corresponds to the number of particles in that angular orientation. **c.** Directional Fourier Shell Correlation (FSC) plot representing 3D resolution anisotropy in the reconstructed map, with the red line showing the global FSC, green dashed lines correspond to ±1 standard deviation from mean of directional resolutions, and the blue histograms correspond to percentage of directional resolution over the 3D FSC. **d.** EM density quality of guide-target duplex. **e.** Individual AtAgo10 domains fit into the EM density, EM density shown in mesh; molecular models (colored as in Fig. 1) shown in cartoon representation with side chains shown as sticks. **f.** Close-up view of density surrounding the scissile phosphate (yellow).

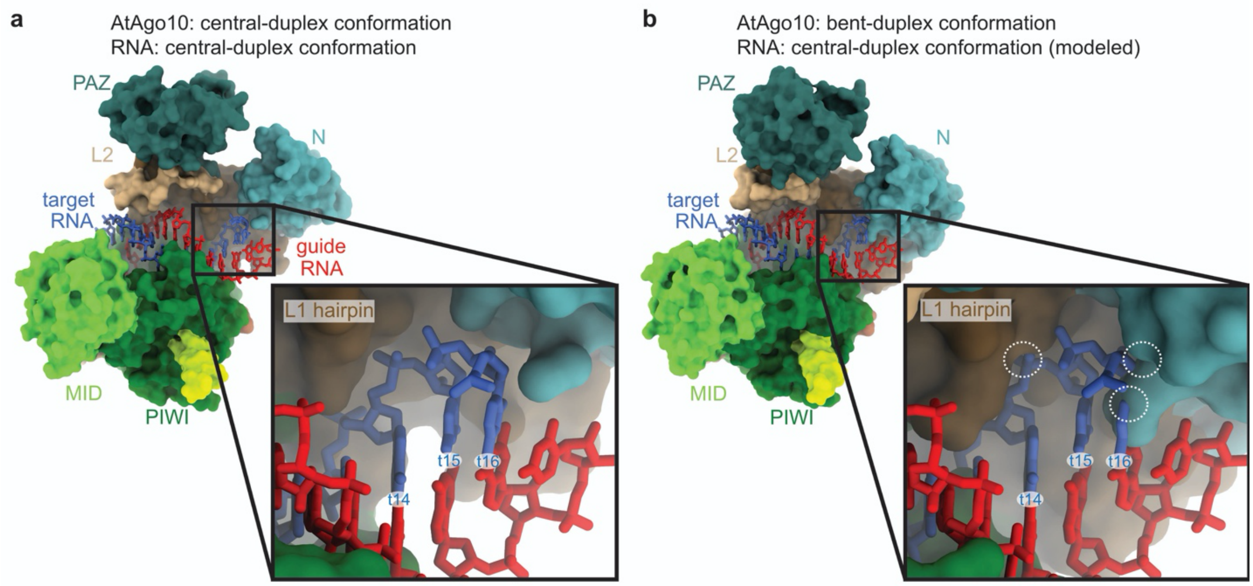

**Fig. S6. Bent-duplex conformation of AtAgo10 cannot bind a central guide-target duplex with pairing beyond t13.**

**a.** Surface representation of the AtAgo10 central-duplex structure with the guide-target RNA duplex shown as sticks. Inset shows no steric clashes between AtAgo10 and the end of the RNA duplex. b. Surface representation of AtAgo10 in the bent-duplex conformation with a modeled continuous guide-target RNA duplex (taken from the central-duplex structure). Inset shows steric clashes (white dashed circles) between t14/t16 of the modeled RNA and the L1 hairpin/N domain of AtAgo10.

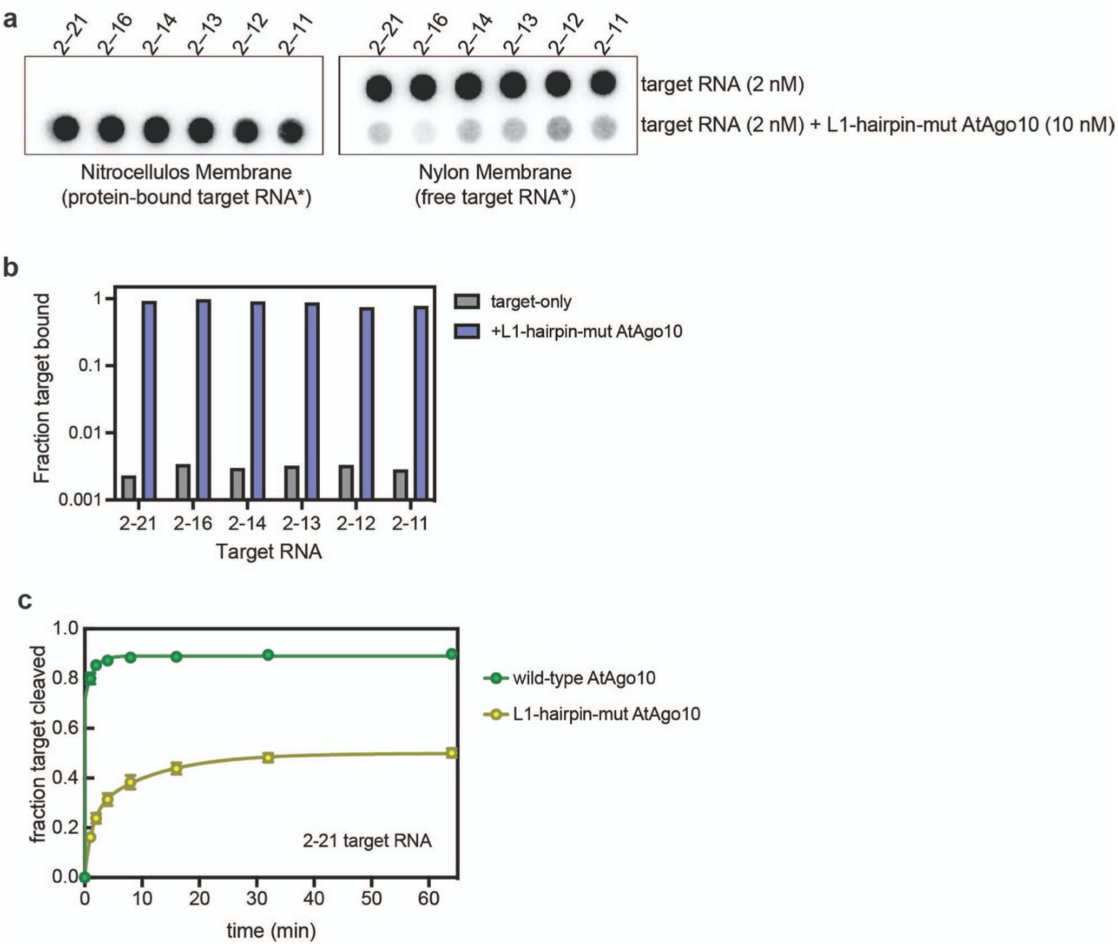

**Fig. S7. L1-hairpin-mut AtAgo10 is impaired in target-slicing but not target-binding**
**a.** Blots from an RNA filter-binding experiment using L1-hairpin-mut AtAgo10 (10 nM) and target RNAs (2 nM) used in this study (shown in Fig. 4B). Complexes were formed with $^{32}$P-labeled target RNAs under conditions used for slicing experiments (except $Mg^{2+}$ was omitted to prevent target cleavage) and passed through a nitrocellulose membrane (to capture protein-RNA complexes) followed by a nylon membrane (to capture RNAs not retained on the nitrocellulose). b. Quantitation of data in panel a. c. Fraction of 2-21 target RNA (2 nM) cleaved by wild-type (10 nM) or L1-hairpin-mut AtAgo10 (10 nM) versus time.

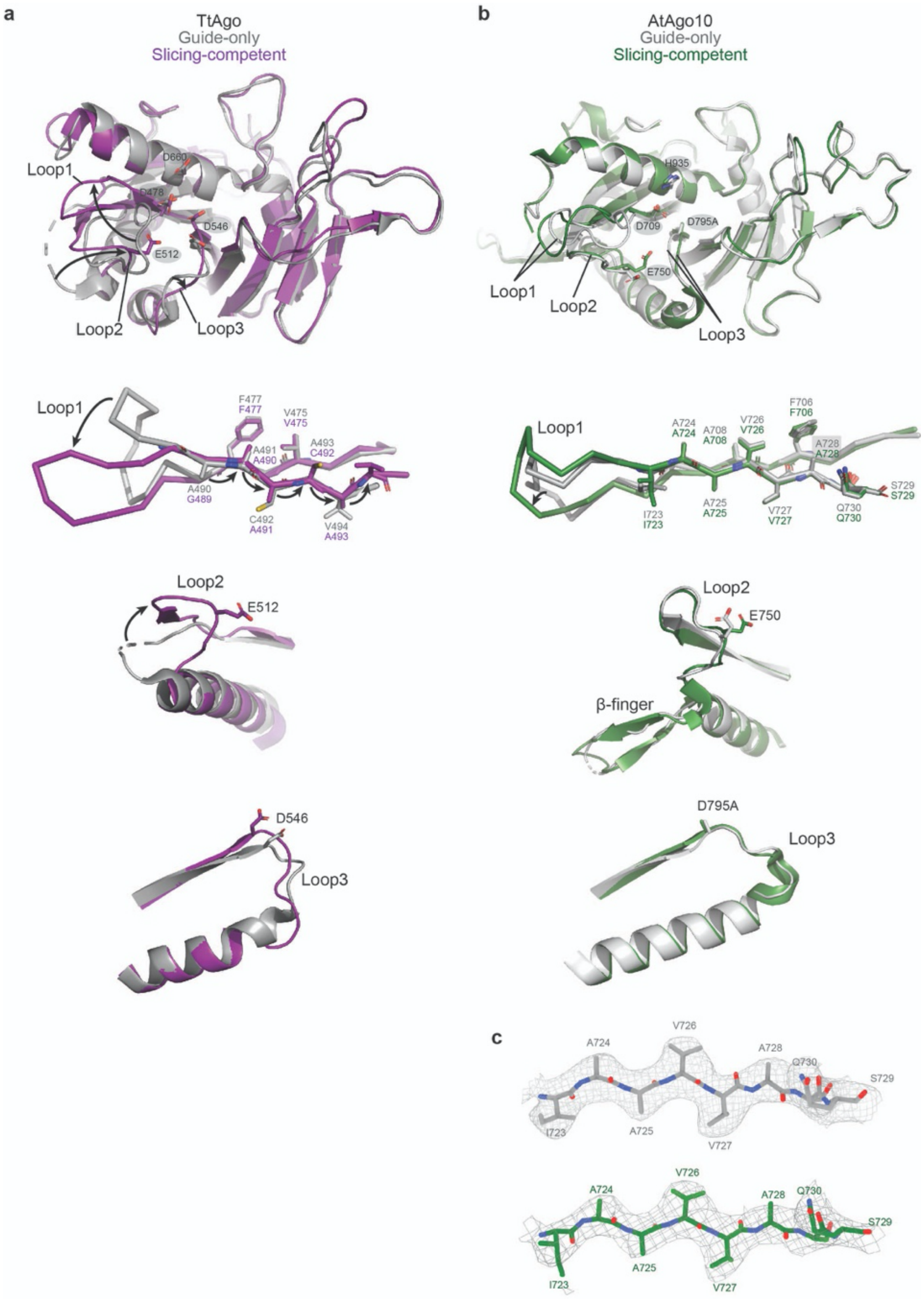

**Fig. S8. Comparison of TtAgo and AtAgo10 PIWI domain conformational changes.**
**a.** Superimposition of TtAgo PIWI domains from guide-only (grey, PDB: 3DLH) and slicing competent (purple, PDB: 4NCB) conformations. Top panel: an overview of the TtAgo PIWI domain with active site residues shown as sticks and labeled. Bottom panel: Detailed views showing the formation of the catalytic conformation involves three rearranged loops and flipping+translation of a ß-strand connected to loop-1 in the TtAgo PIWI domain. Arrows indicate major shifts from the guide-only to the slicing-competent conformation. **b.** Superimposition of the AtAgo10 PIWI

domains from guide-only (grey) and slicing competent (central-duplex) conformations (green) shows relatively few changes and no ß-strand repositioning. **c.** Cryo-EM density map (mesh) around the AtAgo10 β-strand attached to loop1 is shown to illustrate no ß-strand repositioning (grey: guide-only conformation. green: slicing-competent conformation).

**Table S1. Cryo-EM data collection, refinement, and validation statistics.**

| Sample name | AtAgo10-guide RNA | AtAgo10-guide-target (central duplex) | AtAgo10-guide-target (bent duplex) |
|---|---|---|---|
| EMDB ID | EMD-25446 | EMD-25472 | EMD-25482 |
| PDB ID | 7SVA | 7SWF | 7SWQ |
| Microscope | Talos Arctica | Talos Arctica | |
| Detector (Mode) | Gatan K2 Summit, counting mode | Gatan K2 Summit, counting mode | |
| Voltage (kV) | 200 | 200 | |
| Magnification (nominal) | 73,000x | 45,000x | |
| Total electron fluence ($e^{-}/Å^{2}$) | 66.51 | 66.88 | |
| Defocus range (μm) | -0.8 to -1.3 | -1.0 to -1.7 | |
| Pixel size (Å) | 0.566 | 0.91 | |
| Total exposure time (sec) | 11 | 8.4 | |
| Total frames/micrograph | 55 | 42 | |
| Exposure per frame ($e^{-}/Å^{2}$/frame) | 1.21 | 1.59 | |
| Micrographs collected (no.) | 2,656 | 2,049 | |
| Total extracted particles (no.) | 736,209 | 1,935,081 | |
| Particles used for 3D analyses (no.) | 381,087 | 405,690 | 190,349 |
| Final refined particles (no.) | 53,766 | 38,833 | 28,499 |
| Symmetry imposed | C1 | C1 | C1 |
| Map Global Resolution (Å) | 3.26 | 3.79 | 3.79 |
| FSC threshold | 0.143 | 0.143 | 0.143 |
| FSC Sphericity | 0.963 | 0.964 | 0.949 |
| Local resolution range (Å) | 3.0 – 6.0 | 3.5 – 6.5 | 3.5-6.5 |
| Map Sharpening *B* factors ($Å^{2}$) | -115.7 | -70 | -88.3 |
| **Refinement** | | | |
| Refinement package (s) | Phenix | Phenix | Phenix |
| Model composition | | | |
| Non-hydrogen atoms | 6515 | 6873 | 6998 |

| | | | |
|---|---|---|---|
| Protein residues | 792 | 778 | 791 |
| RNA residues | 11 | 34 | 35 |
| $Mg^{2+}$ ions | 1 | 1 | 1 |

| | | | |
|---|---|---|---|
| Model resolution (Å) | | | |
| FSC 0.5<br>FSC 0.143 | 3.4<br>3.1 | 3.9<br>3.6 | 3.9<br>3.6 |
| *B* factors (Å$^2$) | | | |
| Protein | 40.38 | 83.05 | 34.47 |
| Nucleotide | 39.44 | 93.19 | 90.16 |
| Map Correlation Coefficient | | | |
| Global | 0.82 | 0.82 | 0.76 |
| Local | 0.79 | 0.80 | 0.74 |
| R.m.s. deviations | | | |
| Bond lengths (Å) | 0.004 | 0.010 | 0.004 |
| Bond angles (°) | 0.957 | 0.976 | 0.739 |
| **Validation** | | | |
| EMRinger score | 3.41 | 2.89 | 2.76 |
| MolProbity score | 1.85 | 2.31 | 2.19 |
| Clashscore | 8.64 | 15.32 | 12.99 |
| Rotamer outliers (%) | 0.29 | 1.33 | 0.00 |
| Cβ deviations (%) | 0.00 | 0.00 | 0.00 |
| Ramachandran plot | | | |
| Favored (%) | 94.36 | 90.89 | 89.35 |
| Allowed (%) | 5.64 | 9.11 | 10.65 |
| Disallowed (%) | 0.00 | 0.00 | 0.00 |
